# A Nimbolide-Based Kinase Degrader Preferentially Degrades Oncogenic BCR-ABL

**DOI:** 10.1101/2020.04.02.022541

**Authors:** Bingqi Tong, Jessica N. Spradlin, Luiz F.T. Novaes, Erika Zhang, Xirui Hu, Malte Moeller, Scott M. Brittain, Lynn M. McGregor, Jeffrey M. McKenna, John A. Tallarico, Markus Schirle, Thomas J. Maimone, Daniel K. Nomura

## Abstract

Targeted protein degradation (TPD) and proteolysis-targeting chimeras (PROTACs) have arisen as powerful therapeutic modalities for degrading specific protein targets in a proteasome-dependent manner. However, a major limitation to broader TPD applications is the lack of E3 ligase recruiters. Recently, we discovered the natural product nimbolide as a covalent ligand for the E3 ligase RNF114. When linked to the BET family inhibitor JQ1, the resulting heterobifunctional PROTAC molecule was capable of selectively degrading BRD4 in cancer cells. Here, we show the broader utility of nimbolide as an E3 ligase recruiter for TPD applications. We demonstrate that a PROTAC linking nimbolide to the kinase and BCR-ABL fusion oncogene inhibitor dasatinib, BT1, selectively degrades BCR-ABL over c-ABL in leukemia cancer cells, compared to previously reported cereblon or VHL-recruiting BCR-ABL degraders that show opposite selectivity or in some cases inactivity. Further contrasting from cereblon or VHL-recruiting degradation, we show that BT1 treatment not only leads to BCR-ABL degradation, but also stabilizes the endogenous RNF114 substrate and tumor suppressor substrate p21. This leads to additional anti-proliferative effects in leukemia cancer cells beyond those observed with cereblon or VHL-recruiting BCR-ABL PROTACs. Thus, we further establish nimbolide as an additional general E3 ligase recruiter for PROTACs with unique additional benefits for oncology applications. We also further demonstrate the importance of expanding upon the arsenal of E3 ligase recruiters, as such molecules confer differing and unpredictable selectivity for the degradation of neo-substrate proteins.

## Main text

Targeted protein degradation (TPD) and proteolysis-targeting chimeras (PROTACs) are powerful therapeutic paradigms that employ heterobifunctional molecules to recruit an E3 ligase to a protein of interest for polyubiquitination and degradation by the proteasome (Burslem and Crews, 2017; Lai and Crews, 2017). While this technology is a very promising drug discovery paradigm for tackling so far intractable therapeutic targets, and small-molecule PROTACs have entered human clinical trials (Mullard, 2019), a major challenge in the application of this technology is the small number of known E3 ligase recruiters. While there are ∼600 different E3 ligases, many e.g. with different cellular localization, only a few E3 ligase recruiters have been identified and successfully employed, including small-molecule recruiters for cereblon (CRBN), VHL, MDM2, and cIAP (Lai and Crews, 2017; Rape, 2018); amongst these, most reported degraders have used either CRBN or VHL ligands. Additionally, recent reports also suggest resistance to degraders may occur through reprogramming of cellular machinery on the E3 ligase binding side, and there are certain proteins that have been resistant to degradation (Zeng et al., 2020; Zhang et al., 2019a). The discovery of new E3 ligase recruiters is thus necessary to expand the scope of TPD applications.

Recently, chemoproteomic platforms have been used to discover additional recruiters that act through covalent targeting of cysteines on E3 ligases. These covalent recruiters include CCW16 that targets RNF4, SB002 that binds to DCAF16, and nimbolide that targets RNF114. To show proof-of-concept, these new recruiters were linked to JQ1 or to an SLF ligand to show proteasome-dependent degradation of BRD4 or FKBP12, respectively (Spradlin et al., 2019; Ward et al., 2019; Zhang et al., 2019b). While these studies have effectively shown that E3 ligases can be targeted covalently for recruitment and degradation of their target proteins, both FKBP12 and BRD4 are among targets that are considered easily degradable, that is, many PROTACs have been developed that efficiently and robustly degrade these targets in a selective and proteasome-dependent manner.

In prior work employing nimbolide-JQ1 degraders we observed selective degradation of BRD4 over related Bromodomain and extraterminal domain (BET) family members (Spradlin et al., 2019). In this study, we further investigated the broader utility and selectivity of nimbolide-based PROTACs against protein targets that possess additional degradation selectivity challenges, namely human kinases. Among these targets, selective targeting of the oncogenic fusion protein BCR-ABL, a driver of chronic myelogenous leukemia (CML), over c-ABL, an important non-receptor tyrosine kinase involved in numerous cellular processes (Hantschel and Superti-Furga, 2004), have proven to be challenging with studied PROTACs exploiting CRBN or VHL recruitment. Pioneering studies by the Crews lab have shown that BCR-ABL PROTACs, consisting of CRBN or VHL recruiters linked to BCR-ABL ATP binding pocket-targeting kinase inhibitors such as dasatinib and bosutinib or allosteric inhibitors such as GNF-2, showed preferential degradation of c-ABL over modest degradation of BCR-ABL (Burslem et al., 2019; Lai et al., 2016). Notably, this work has spurred development of various degradation strategies targeting BCR-ABL (Demizu et al., 2016; Shibata et al., 2017, 2019; Shimokawa et al., 2017; Zhao et al., 2019). In this study, we aimed to determine whether our recently discovered covalent RNF114 recruiter nimbolide could be used to kinase-targeting PROTACs, and if so, whether differential selectivity in degrading BCR-ABL, compared to similar BCR-ABL PROTACs employing VHL or CRBN could be attained.

The scope of targets amenable to nimbolide-based, RNF114-mediated protein degradation has without question been hampered by the extreme cost of nimbolide (2020 Sigma price = $77,000 USD/gram), yet commercial neem leaf powder (*azadirachta indica* extract) remains a popular and inexpensive health supplement containing varying amounts of nimbolide. Using a newly devised extraction protocol (see Supporting Information), we were able to obtain gram quantities of analytically pure nimbolide from a single 1 lb. can of Organic Veda™ neem leaf extract (2020 amazon.com price = $14.99 USD) without the need for HPLC purification **(Figure 1)**. Importantly, this process opens the door to the synthetic and chemical biology communities to manipulate this complex triterpene scaffold for biological and especially TPD applications.

**Figure 1.**
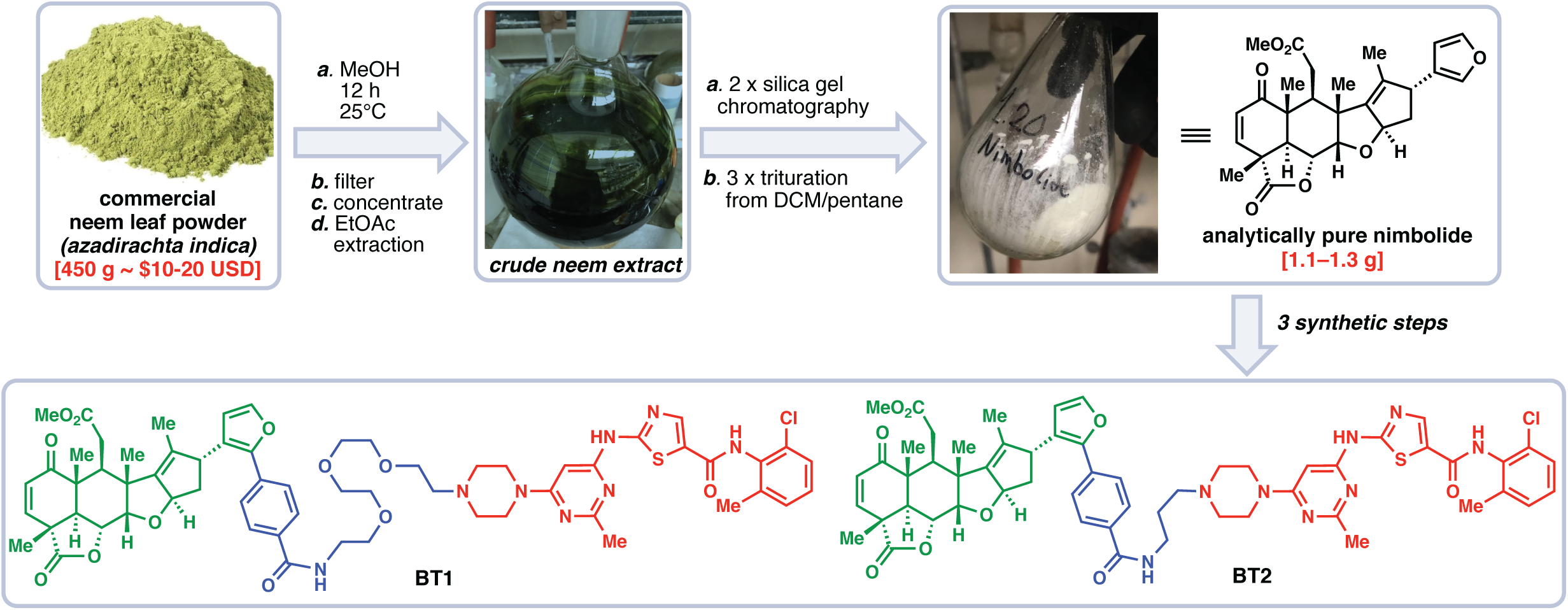
Extracting nimbolide from neem and nimbolide-based BCR-ABL degraders. Efficient method for extracting gram quantities of nimbolide from neem. This procedure is described in detail in Supporting Information. Structures of the nimbolide-based BCR-ABL degraders BT1 and BT2 are shown.

With access to significant quantities of nimbolide, we synthesized two nimbolide-based degraders linked to the tyrosine kinase inhibitor dasatinib employing both short alkyl (see BT2) and longer PEG-based (see BT1) linkers in analogy to previous work (**Figure 1**, see Supporting Information for synthetic details) (Spradlin et al., 2019). Treatment of K562 leukemia cells expressing the fusion oncogene BCR-ABL for 24 h with BT1 and BT2 led to loss of both BCR-ABL and c-ABL, with a more pronounced effect observed with BT1 **(Figure 2a)**. While our prior work on BRD4 degradation with Nimbolide/JQ-1-based PROTACS had found shorter linkers to be superior, the deeper binding pocket of kinase ligands may explain the improved performance of BT1 in this context (Spradlin et al., 2019). Interestingly, both BT1 and BT2 showed more degradation of BCR-ABL than c-ABL, demonstrating opposite preference to Crews’ findings **(Figure 2a)**. We show that phosphorylated CRKL, downstream of c-ABL signaling was inhibited by both BT1 and BT2, indicating that BT1 and BT2 sufficiently engaged ABL in cells **(Figure 2a).** While performing rescue studies with proteasome inhibitors was challenging due to the toxicity of proteasome inhibitors at 24 h in this cell line, the degradation of BCR-ABL or c-ABL with BT1 treatment was not observed with nimbolide or dasatinib treatment, nor with co-treatment of both compounds **(Figure 2b).**

**Figure 2.**
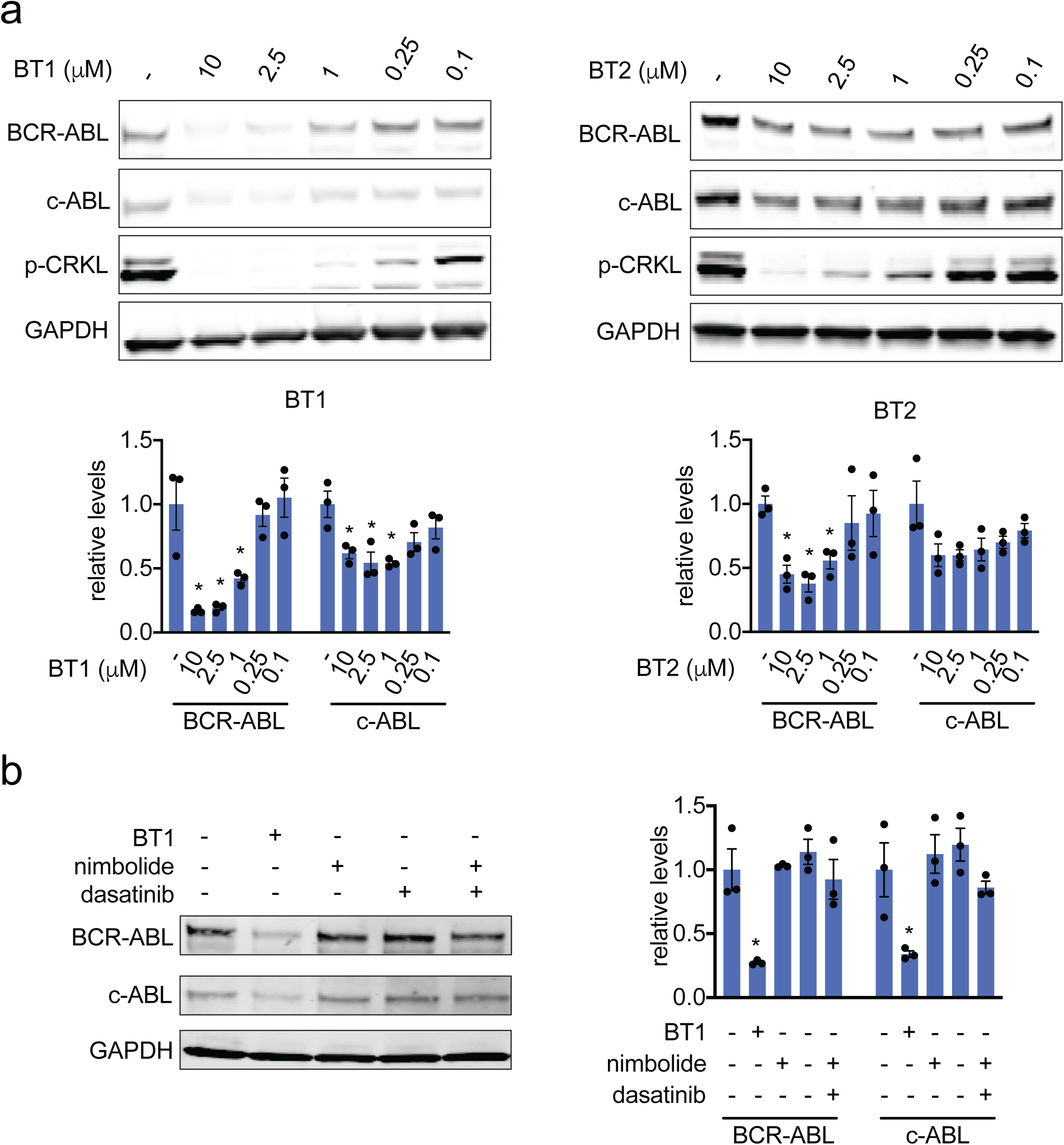
Effects of Nimbolide-based BCR-ABL degraders on BCR-ABL and c-ABL. **(a)** BCR-ABL, c-ABL, phosphorylated CRKL, and loading control GAPDH levels with DMSO vehicle, BT1, and BT2 treatment in K562 cells for 24 h, assessed by Western blotting and quantified below in bar graphs by densitometry and normalization to GAPDH. **(b)** BCR-ABL, c-ABL, and GAPDH loading control levels in K562 cells treated with BT1 (1 μM), nimbolide (1 μM), or dasatinib (1 μM) for 24 h, assessed by Western blotting and quantified by densitometry normalized to GAPDH. Blots are representative of n=3 biological replicates/group. Quantified data show individual replicate values, average, and sem. Statistical significance is expressed as *p<0.05 compared to vehicle-treated control for each group.

We next performed a more detailed time-course analysis comparing our BT1 degrader with previously reported CRBN and VHL-recruiting dasatinib degraders of related linker length and composition **(Figure 3a-3c, Supporting Figure S1)**. Consistent with previous reports, the CRBN-dasatinib PROTAC showed preferential and faster degradation of c-ABL compared to BCR-ABL **(Figure 3a)**. We also observed no significant degradation of BCR-ABL during the 24 h time-course with >50 % of degradation of c-ABL when employing the VHL-dasatinib degrader, consistent with Crews’ findings **(Figure 3b)**. In contrast, BT1 showed faster and preferential degradation of BCR-ABL over c-ABL at every time-point tested **(Figure 3c)**.

**Figure 3.**
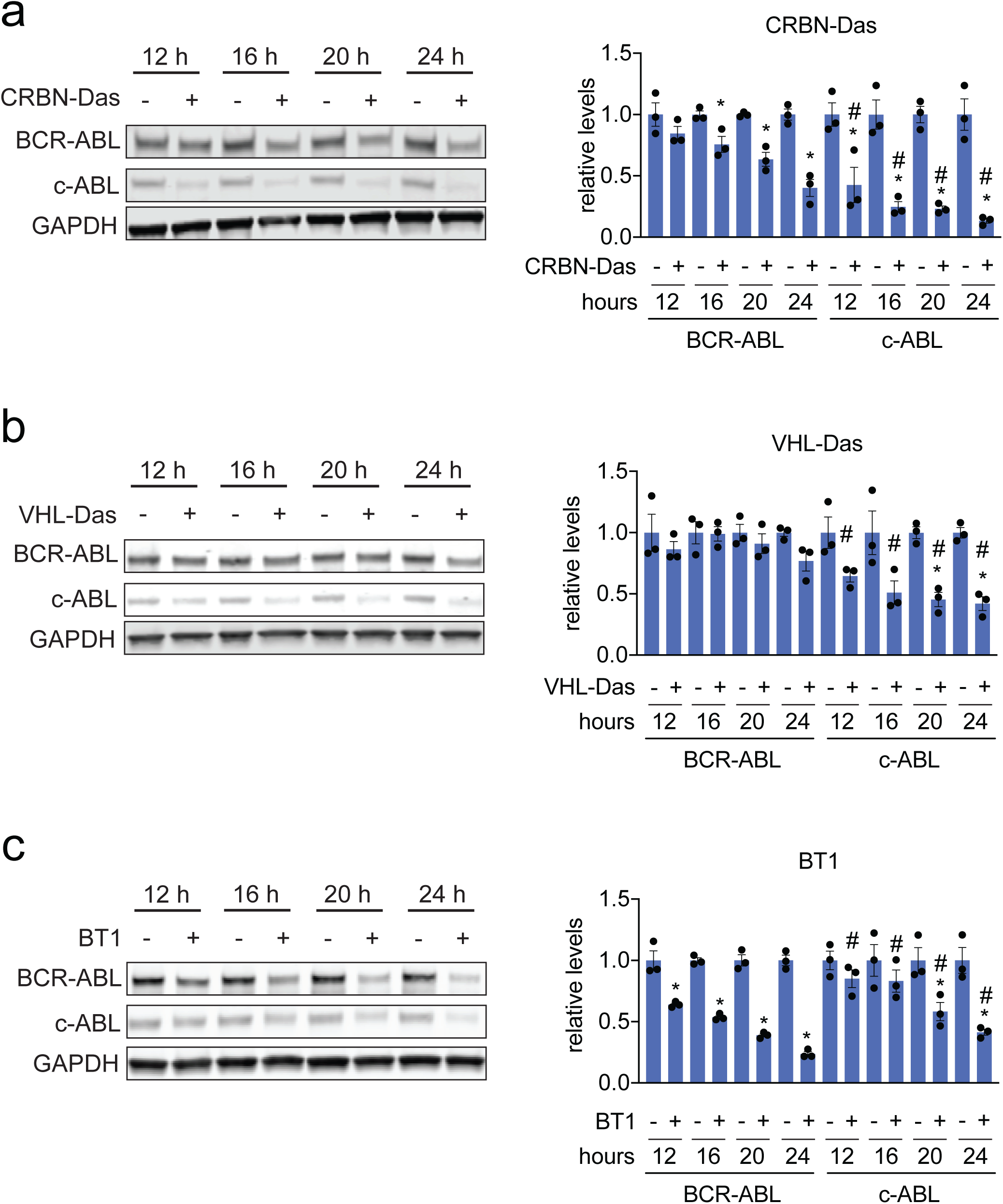
Rate of BCR-ABL versus c-ABL degradation with RNF114, CRBN, or VHL-based BCR-ABL degraders. **(a, b, c)** BCR-ABL, c-ABL, and loading control GAPDH levels with K562 cell treatment with DMSO vehicle or CRBN (thalidomide)-, VHL (VHL ligand), or RNF114 (nimbolide)-based (BT1) dasatinib BCR-ABL degraders at 2.5 μM for 12, 16, 20, or 24 h, assessed by Western blotting and quantified by densitometry and normalized to GADPH loading control. Structures of all three degraders are show in **Supporting Figure S1.** Blots are representative of n=3 biological replicates/group. Quantified data show individual replicate values, average, and sem. Statistical significance is expressed as *p<0.05 compared to vehicle-treated control for each group, #p<0.05 compared to the corresponding BCR-ABL treatment comparisons with the individual degraders.

We had previously reported that nimbolide disrupts endogenous RNF114 substrate recognition through targeting an N-terminal cysteine (Cys 8), leading to accumulation of endogenous RNF114 substrates such as the tumor suppressors such as CDKN1A (p21) and CDKN1C (p57) which in turn results in impaired breast cancer cell viability. Our previously described nimbolide-JQ1 BRD4 degrader showed selective degradation of BRD4, but also enhanced levels of p21 and p57 in breast cancer cells. We demonstrate here that BT1 treatment also led to greatly elevated levels of CDKN1A (p21) levels in K562 cells—an effect that was not observed with dasatinib, CRBN-dasatinib, or VHL-dasatinib treatments **(Figure 4a, 4b).** These results indicate that a nimbolide-based BCR-ABL degrader may possess additional therapeutic properties beyond dasatinib alone. While dasatinib impaired K562 cell proliferation at much lower concentrations compared to BT1 likely due to better cell permeability of dasatinib compared to BT1, we observed significantly greater impairments in cell proliferation with higher concentrations of BT1 from 1 to 5 μM compared to treatment with dasatinib alone, nimbolide alone, or even dasatinib and nimbolide together **(Figure 4c).** While we do not yet understand why BT1 shows more efficacy at these higher concentrations than with dasatinib and nimbolide co-treatment, we postulate that the observed heightened viability impairments observed with BT1 are due to the additional anti-cancer effects from nimbolide targeting of RNF114. These effects are likely observed only at the higher concentrations due to the poorer anti-proliferative potency of nimbolide compared to dasatinib **(Figure 4c).** We also observed significantly greater anti-proliferative effects with BT1 from 1 to 5 μM, compared to CRBN-dasatinib and VHL-dasatinib PROTACs **(Figure 4d)**. Additionally, while CRBN-dasatinib potently ablates the known dasatinib target Bruton’s tyrosine kinase (BTK), and VHL-dasatinib shows robust, but diminished degradation, substantially lower levels of degradation for these kinases is noted with BT1 treatment **(Figure 4a, 4b)**. These findings further highlight the emerging complexities inherent to kinase degradation (Bondeson et al., 2018; Brand et al., 2019; Cromm et al., 2018; Huang et al., 2018; Jaime-Figueroa et al., 2020; Li et al., 2020; Olson et al., 2018; Powell et al., 2018; Tovell et al., 2019; You et al., 2020).

**Figure 4.**
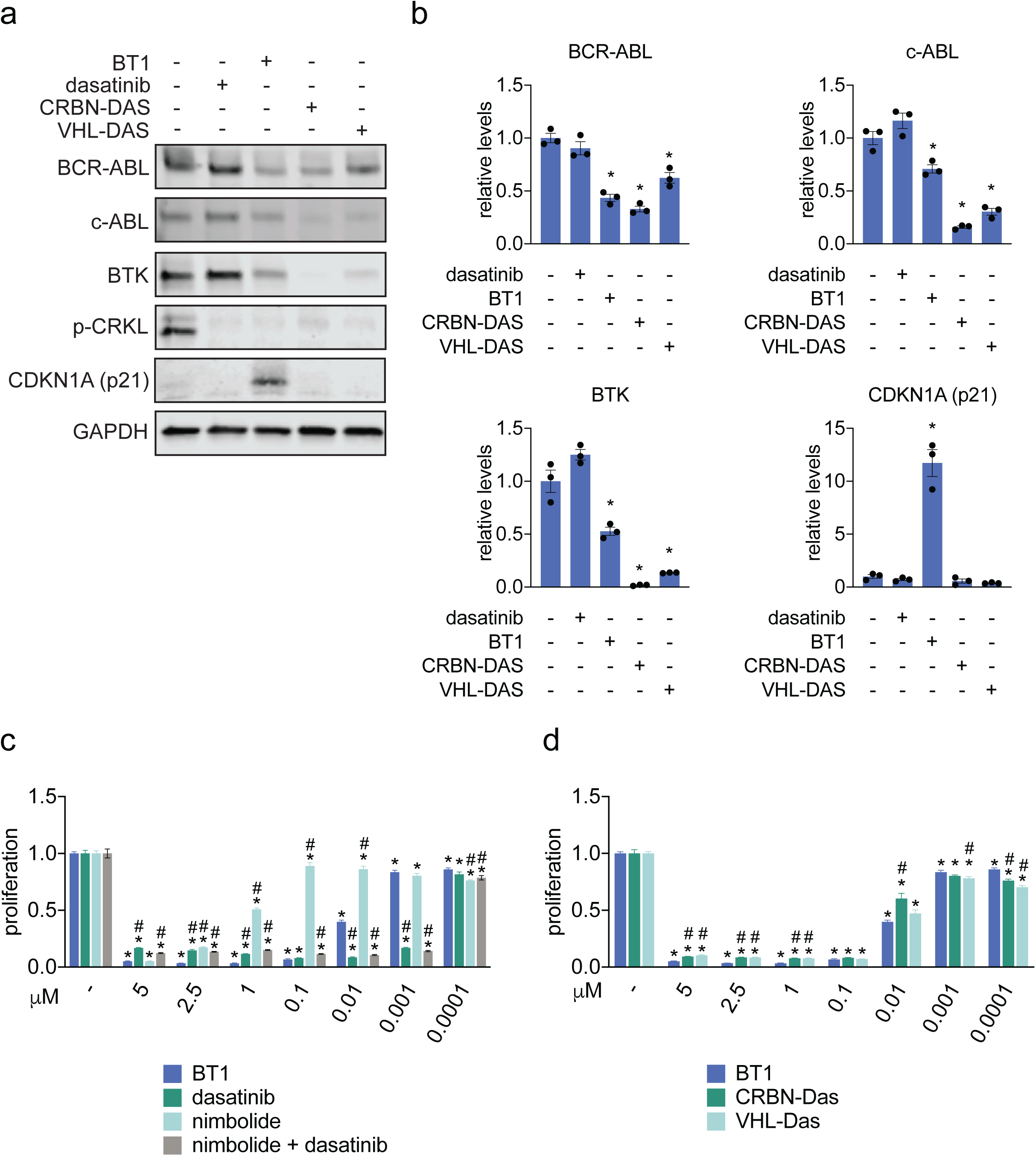
Comparing RNF114, CRBN, or VHL-recruiting BCR-ABL degraders. **(a, b)** BCR-ABL, c-ABL, BTK, phosphorylated CRKL, CDKN1A (p21), and loading control GAPDH levels in K562 cells treated with DMSO vehicle or dasatinib (2.5 μM), BT1 (2.5 μM), CRBN-DAS (2.5 μM), and VHL-DAS (2.5 μM) for 24 h, assessed by Western blotting and quantified in **(b)** by densitometry and normalized to GADPH loading control. **(c, d)** Cell proliferation of K562 cells treated with DMSO vehicle, BT1, dasatinib, nimbolide, nimbolide and dasatinib combined, CRBN-dasatinib, or VHL-dasatinib for 48 h. Blots are representative of n=3 biological replicates/group. Quantified data show individual replicate values, average, and sem. Statistical significance is expressed as *p<0.05 compared to vehicle-treated control for each group, #p<0.05 compared to BT1 treatment group.

Collectively, we demonstrate that covalent E3 ligase recruiters can be used to degrade kinases in TPD applications, and that a nimbolide-based BCR-ABL degrader shows unique degradation specificity profiles compared with initially reported cereblon- or VHL-recruiting degraders. The propensity of RNF114-recruiting nimbolide-based degraders to degrade the oncogenic protein form more selectively is particularly notable. Additionally, nimbolide-based degraders may also possess additional anti-cancer effects through heightening the levels of tumor-suppressors such as p21. Our results underscore the importance of discovering more E3 ligase recruiters for better tuning the specificity and effects of future clinical candidates and drugs arising from TPD-based approaches. Finally, with ready access to supplies of nimbolide, a study of the scope of RNF114-mediated protein degradation can begin in earnest.

## Acknowledgement

We thank the members of the Nomura Research Group, the Maimone Research Group, and Novartis Institutes for BioMedical Research for critical reading of the manuscript. This work was supported by Novartis Institutes for BioMedical Research and the Novartis-Berkeley Center for Proteomics and Chemistry Technologies (NB-CPACT) for all listed authors. This work was also supported by the Nomura Research Group and the Mark Foundation for Cancer Research ASPIRE Award for DKN, JNS. This work was also supported by grants from the National Institutes of Health (R01CA240981 for DKN, TJM, BT, JNS).

## Author Contributions

DKN, TJM conceived the project and wrote the paper. BT, JNS, SMB, JAT, JMK, LM, MS, TJM, DKN provided intellectual contributions and insights into project direction and designed the experiments. BT, JNS, DKN, SMB, TJM performed experiments and analyzed data. All authors edited the paper.

## Competing Financial Interests Statement

SMB, JAT, JMK, LM, MS are employees of Novartis Institutes for BioMedical Research. This study was funded by the Novartis Institutes for BioMedical Research and the Novartis-Berkeley Center for Proteomics and Chemistry Technologies. DKN is a co-founder, shareholder, and adviser for Artris Therapeutics and Frontier Medicines.

## Online Methods

### Materials

Primary antibodies to ABL (Santa Cruz Biochemicals, 24-11), BTK (Cell Signaling Technologies, D3H5), GAPDH (Proteintech Group Inc., 60004-1-Ig), phosphorylated CRKL Y207 (Cell Signaling Technologies, 3181) and p21 (Cell Signaling Technology, 12D1) were obtained from commercial sources and dilutions were prepared according to manufacturer recommendations. Anti-rabbit and anti-mouse seconadary antibodies were purchased from Licor (IRDye 800CW Goat anti-Rabbit IgG Secondary Antibody and IRDye 700CW Goat anti-Mouse IgG Secondary Antibody).

### Isolation of nimbolide and synthesis of BT1 and BT2

Isolation of nimbolide and synthesis of BT1 and BT2 are described in Supporting Information.

### Cell Culture

K562 cells were obtained from the UCB Cell Culture Facility and were cultured in Iscove’s Modified Dulbecco’s Medium (IMDM) containing 10% (v/v) fetal bovine serum (FBS), maintained at 37 °C with 5% CO2.

### Proliferation Assays

Cell proliferation assays were performed using WST8 reagent (APExBio, CCK-8) following manufacturer’s recommendations. K562 cells were seeded at a density of 100,000 cell/mL in a volume of 100 µL (10,000 cells per well). The next day cells were treated with an additional 25 µL of media containing a 1:200 dilution of 1,000X compound stock in DMSO. After the corresponding incubation period 10 µL of WST8 reagent were added directly to the growth media and plates were returned to the 37 °C incubator for 90 min. After incubation absorbance at 450 nm was read using a VersaMax plate reader (Molecular Devices). WST8 reagent was also added to wells containing only media which were used for background subtraction during analysis. Relative proliferation was calculated as the ratio of Abs(450) of the treated wells to the vehicle treated control wells on the same plate.

### Western Blotting

K562 cells were seeded for treatment at a density of 500,000 cells/mL, treated with compounds dissolved in DMSO and harvested after the appropriate treatment time by centrifugation at 1,000xg for 5 min at 4 °C. Cell pellets were washed with 500 µL phosphate buffered saline and lysed in 75-100 µL Radioimmunoprecipitation assay buffer (RIPA buffer) with protease inhibitor cocktail (Thermo Fisher Scientific, Pierce Protease Inhibitor Mini Tablets, EDTA-free). Cellular debris was pelleted by spinning at maximum speed for 10 minutes at 4 °C. Supernatant was transferred and total protein was normalized by Pierce BCA Protein Assay. Samples were denatured by addition of 4X Laemmli’s Loading dye and 25-50 µg of protein was loaded onto 4-20% TGX Precast gels (BioRad). After gel electrophoresis proteins were transferred to a nitrocellulose membrane using semi-dry transfer on a Trans-Blot Turbo (BioRad) over 10 min. The membrane was then incubated for 1 hour in 5% bovine serum albumin (BSA) in tris-buffered saline containing Tween 20 (TBST) before being incubated with the correct primary antibody overnight at 4 °C. The membranes were washed in TBST before a 1 hour room temperature incubation with secondary antibodies. After a final set of washes blots were imaged on a LiCor CLX imager and band intensitities were quantified using ImageJ software.

### Data Availability Statement

The datasets generated during and/or analyzed during the current study are available from the corresponding author on reasonable request.

## References

Bondeson, D.P., Smith, B.E., Burslem, G.M., Buhimschi, A.D., Hines, J., Jaime-Figueroa, S., Wang, J., Hamman, B.D., Ishchenko, A., and Crews, C.M. (2018). Lessons in PROTAC Design from Selective Degradation with a Promiscuous Warhead. Cell Chem. Biol. 25, 78-87.e5.

Brand, M., Jiang, B., Bauer, S., Donovan, K.A., Liang, Y., Wang, E.S., Nowak, R.P., Yuan, J.C., Zhang, T., Kwiatkowski, N., et al. (2019). Homolog-Selective Degradation as a Strategy to Probe the Function of CDK6 in AML. Cell Chem. Biol. 26, 300-306.e9.

Burslem, G.M., and Crews, C.M. (2017). Small-Molecule Modulation of Protein Homeostasis. Chem. Rev. 117, 11269–11301.

Burslem, G.M., Schultz, A.R., Bondeson, D.P., Eide, C.A., Savage Stevens, S.L., Druker, B.J., and Crews, C.M. (2019). Targeting BCR-ABL1 in Chronic Myeloid Leukemia by PROTAC-Mediated Targeted Protein Degradation. Cancer Res. 79, 4744–4753.

Cromm, P.M., Samarasinghe, K.T.G., Hines, J., and Crews, C.M. (2018). Addressing Kinase-Independent Functions of Fak via PROTAC-Mediated Degradation. J. Am. Chem. Soc. 140, 17019–17026.

Demizu, Y., Shibata, N., Hattori, T., Ohoka, N., Motoi, H., Misawa, T., Shoda, T., Naito, M., and Kurihara, M. (2016). Development of BCR-ABL degradation inducers via the conjugation of an imatinib derivative and a cIAP1 ligand. Bioorg. Med. Chem. Lett. 26, 4865–4869.

Hantschel, O., and Superti-Furga, G. (2004). Regulation of the c-Abl and Bcr-Abl tyrosine kinases. Nat. Rev. Mol. Cell Biol. 5, 33–44.

Huang, H.-T., Dobrovolsky, D., Paulk, J., Yang, G., Weisberg, E.L., Doctor, Z.M., Buckley, D.L., Cho, J.-H., Ko, E., Jang, J., et al. (2018). A Chemoproteomic Approach to Query the Degradable Kinome Using a Multi-kinase Degrader. Cell Chem. Biol. 25, 88-99.e6.

Jaime-Figueroa, S., Buhimschi, A.D., Toure, M., Hines, J., and Crews, C.M. (2020). Design, synthesis and biological evaluation of Proteolysis Targeting Chimeras (PROTACs) as a BTK degraders with improved pharmacokinetic properties. Bioorg. Med. Chem. Lett. 30, 126877.

Lai, A.C., and Crews, C.M. (2017). Induced protein degradation: an emerging drug discovery paradigm. Nat. Rev. Drug Discov. 16, 101–114.

Lai, A.C., Toure, M., Hellerschmied, D., Salami, J., Jaime-Figueroa, S., Ko, E., Hines, J., and Crews, C.M. (2016). Modular PROTAC Design for the Degradation of Oncogenic BCR-ABL. Angew. Chem. Int. Ed Engl. 55, 807–810.

Li, Z., Pinch, B.J., Olson, C.M., Donovan, K.A., Nowak, R.P., Mills, C.E., Scott, D.A., Doctor, Z.M., Eleuteri, N.A., Chung, M., et al. (2020). Development and Characterization of a Wee1 Kinase Degrader. Cell Chem. Biol. 27, 57-65.e9.

Mullard, A. (2019). First targeted protein degrader hits the clinic. Nat. Rev. Drug Discov. 18, 237–239.

Olson, C.M., Jiang, B., Erb, M.A., Liang, Y., Doctor, Z.M., Zhang, Z., Zhang, T., Kwiatkowski, N., Boukhali, M., Green, J.L., et al. (2018). Pharmacological perturbation of CDK9 using selective CDK9 inhibition or degradation. Nat. Chem. Biol. 14, 163–170.

Powell, C.E., Gao, Y., Tan, L., Donovan, K.A., Nowak, R.P., Loehr, A., Bahcall, M., Fischer, E.S., Jänne, P.A., George, R.E., et al. (2018). Chemically Induced Degradation of Anaplastic Lymphoma Kinase (ALK). J. Med. Chem. 61, 4249–4255.

Rape, M. (2018). Ubiquitylation at the crossroads of development and disease. Nat. Rev. Mol. Cell Biol. 19, 59–70.

Shibata, N., Miyamoto, N., Nagai, K., Shimokawa, K., Sameshima, T., Ohoka, N., Hattori, T., Imaeda, Y., Nara, H., Cho, N., et al. (2017). Development of protein degradation inducers of oncogenic BCR-ABL protein by conjugation of ABL kinase inhibitors and IAP ligands. Cancer Sci. 108, 1657–1666.

Shibata, N., Ohoka, N., Hattori, T., and Naito, M. (2019). Development of a Potent Protein Degrader against Oncogenic BCR-ABL Protein. Chem. Pharm. Bull. (Tokyo) 67, 165–172.

Shimokawa, K., Shibata, N., Sameshima, T., Miyamoto, N., Ujikawa, O., Nara, H., Ohoka, N., Hattori, T., Cho, N., and Naito, M. (2017). Targeting the Allosteric Site of Oncoprotein BCR-ABL as an Alternative Strategy for Effective Target Protein Degradation. ACS Med. Chem. Lett. 8, 1042–1047.

Spradlin, J.N., Hu, X., Ward, C.C., Brittain, S.M., Jones, M.D., Ou, L., To, M., Proudfoot, A., Ornelas, E., Woldegiorgis, M., et al. (2019). Harnessing the anti-cancer natural product nimbolide for targeted protein degradation. Nat. Chem. Biol. 15, 747–755.

Tovell, H., Testa, A., Zhou, H., Shpiro, N., Crafter, C., Ciulli, A., and Alessi, D.R. (2019). Design and Characterization of SGK3-PROTAC1, an Isoform Specific SGK3 Kinase PROTAC Degrader. ACS Chem. Biol. 14, 2024–2034.

Ward, C.C., Kleinman, J.I., Brittain, S.M., Lee, P.S., Chung, C.Y.S., Kim, K., Petri, Y., Thomas, J.R., Tallarico, J.A., McKenna, J.M., et al. (2019). Covalent Ligand Screening Uncovers a RNF4 E3 Ligase Recruiter for Targeted Protein Degradation Applications. ACS Chem. Biol. 14, 2430–2440.

You, I., Erickson, E.C., Donovan, K.A., Eleuteri, N.A., Fischer, E.S., Gray, N.S., and Toker, A. (2020). Discovery of an AKT Degrader with Prolonged Inhibition of Downstream Signaling. Cell Chem. Biol. 27, 66-73.e7.

Zeng, M., Xiong, Y., Safaee, N., Nowak, R.P., Donovan, K.A., Yuan, C.J., Nabet, B., Gero, T.W., Feru, F., Li, L., et al. (2020). Exploring Targeted Degradation Strategy for Oncogenic KRASG12C. Cell Chem. Biol. 27, 19-31.e6.

Zhang, L., Riley-Gillis, B., Vijay, P., and Shen, Y. (2019a). Acquired Resistance to BET-PROTACs(Proteolysis Targeting Chimeras) Caused by Genomic Alterations in Core Components of E3 ligase Complexes. Mol. Cancer Ther.

Zhang, X., Crowley, V.M., Wucherpfennig, T.G., Dix, M.M., and Cravatt, B.F. (2019b). Electrophilic PROTACs that degrade nuclear proteins by engaging DCAF16. Nat. Chem. Biol. 15, 737–746.

Zhao, Q., Ren, C., Liu, L., Chen, J., Shao, Y., Sun, N., Sun, R., Kong, Y., Ding, X., Zhang, X., et al. (2019). Discovery of SIAIS178 as an Effective BCR-ABL Degrader by Recruiting Von Hippel-Lindau (VHL) E3 Ubiquitin Ligase. J. Med. Chem. 62, 9281–9298.

