## Supporting Information for "A Nimbolide-Based Kinase Degrader Preferentially Degrades Oncogenic BCR-ABL"

**General Procedures**

Unlessotherwise noted, all reactions were performed in flame-dried glassware under positive pressure of nitrogen or argon. Air- and moisture-sensitive liquids were transferred via syringe. Dry dichloromethane, *N*,*N*-Dimethylformamide and tetrahydrofuran were obtained by passing these previously degassed solvents through activated alumina columns. Organic Neem leaf powder from Organic Veda$^{TM}$ (16 Oz. can) was purchased through Amazon.com (https://www.amazon.com/Organic-Neem-Leaf-Powder-Supplement/dp/B00PYQ6K8G/ref=sr_1_5?keywords=organic%2Bveda%2Bneem%2Bpowder&qid=1583032774&sr=8-5&th=1). *N*-(2-Chloro-6-methylphenyl)-2-[(6-chloro-2-methyl-4-pyrimidinyl)amino]-5-thiazolecarboxamide was purchased from Combi-Blocks Inc. All reagents were used as received from commercial sources, unless stated otherwise. Reactions were monitored by thin layer chromatography (TLC) on TLC silica gel 60 $F_{254}$ glass plates (EMD Millipore) and visualized by UV irradiation and staining with *p*-anisaldehyde, phosphomolybdic acid, or Ninhydrin. Volatile solvents were removed under reduced pressure using a rotary evaporator. Flash column chromatography was performed using Silicycle F60 silica gel (60Å, 230-400 mesh, 40-63 μm). Proton nuclear magnetic resonance ($^{1}H$ NMR) and carbon nuclear magnetic resonance ($^{13}C$ NMR) spectra were recorded on Bruker AV-600 and AV-700 spectrometers operating at 600 and 700 MHz for $^{1}H$ NMR, and 151 and 176 MHz for $^{13}C$ NMR. Chemical shifts are reported in parts per million (ppm) with respect to the residual solvent signal $CDCl_3$ ($^{1}H$ NMR: δ 7.26; $^{13}C$ NMR: δ 77.16), $CD_2Cl_2$ ($^{1}H$ NMR: δ 5.32; $^{13}C$ NMR: δ 53.84), $(CD_3)_2CO$ ($^{1}H$ NMR: δ 2.05; $^{13}C$ NMR: δ 29.84), $(CD_3)_2SO$ ($^{1}H$ NMR: δ 2.50; $^{13}C$ NMR: δ 39.52). Peak multiplicities are reported as follows: *s* = singlet, *d* = doublet, *t* = triplet, *dd* = doublet of doublets, *tt* = triplet of triplets, *m* = multiplet, *br* = broad signal. IR spectra were recorded on a Bruker Vertex 80 FTIR spectrometer. High-resolution mass spectra (HRMS) were obtained by the QB3/chemistry mass spectrometry facility at the University of California, Berkeley using a Thermo LTQ-FT mass spectrometer with electrospray ionization (ESI) technique. Microwave reactions were performed in a Biotage Initiator EXP microwave reactor.

**Supplementary Procedure 1.** Isolation of nimbolide from commercial neem leaf powder

**Initial Extraction.** To a 3 L round bottom flask containing 1 lb. (~450 g) of Organic Veda™ neem leaf powder and a large stir bar was added 1.5 L of anhydrous methanol and the resulting suspension was stirred for 12 hours at room temperature in a fumehood with the lights off. The mixture was then filtered through a short pad of Celite® (typically in 5 portions for faster filtering, filtration takes >1 hour) and the solid was washed with additional methanol until the Celite® layer was white colored (see SI-figure 1B). The filtrate was concentrated *in vacuo* at 30 °C to provide a dark green oil which was partitioned between EtOAc (500 mL) and water (300 mL). The phases were separated, and the aqueous layer was extracted with additional EtOAc (250 mL). The combined organic layers were washed with a mixture of saturated aqueous $NaHCO_3$ and $Na_2S_2O_3$ (9:1 v/v, 250 mL), dried over $Na_2SO_4$ and concentrated *in vacuo* to afford a dark green oil. Thin layer chromatographic (TLC) analysis of this mixture is shown in SI-Figure 1D.

**A** **B** **C** **D**

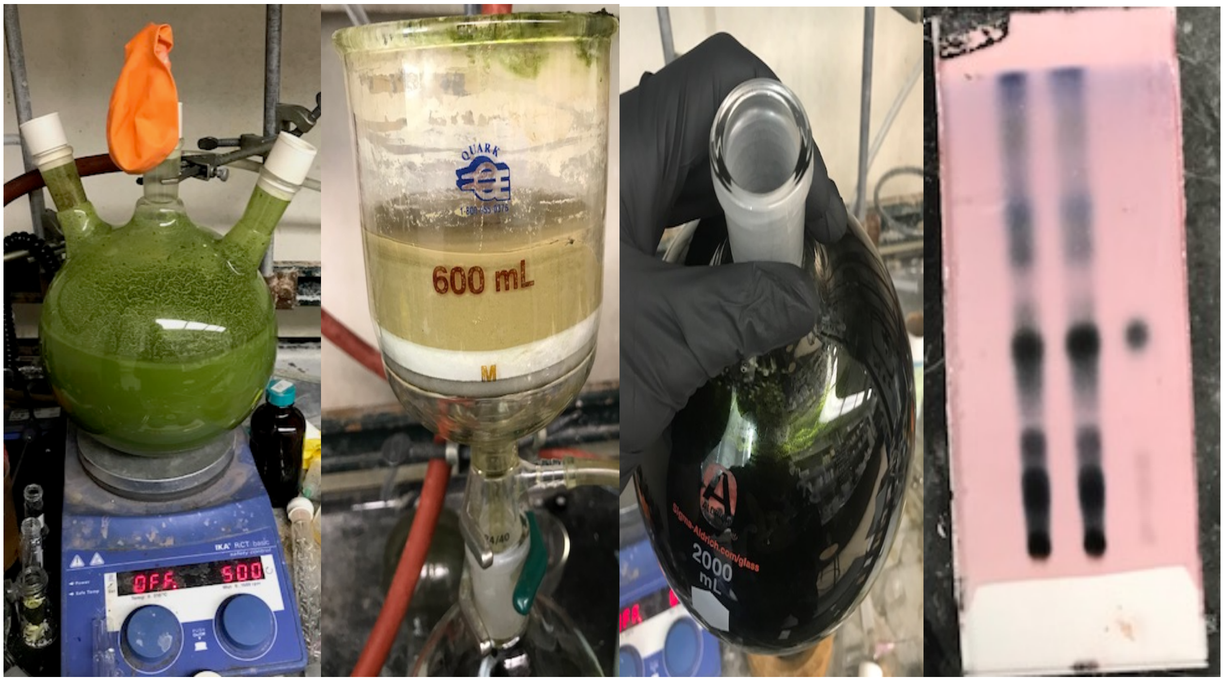


**SI-Figure 1.** Initial extraction step of nimbolide from neem leaf powder. **A**) extraction set-up. **B**) completed filtration through celite. **C**) crude residue obtained. **D**) Thin layer chromatographic analysis (50:50 EtOAc/Hexane, *p*-anisaldehyde stain) of the crude residue. Pure nimbolide is shown in the far-right lane, unpurified mixture in the left lane, and a co-spot in the center lane.

**First Column Chromatography.** The dark green oil was purified by silica gel flash column chromatography according to the following procedure. A column of silica gel (7 cm diameter X 25 cm height) was packed and flushed with hexanes/EtOAc/$Et_3N$ (75:25:1 v/v/v). Next, 100 mL of hexanes was loaded on the top and a solution of the green residue in 50 mL of EtOAc was loaded mixing with the neat hexanes layer. The column was eluted using the following gradient: Hexanes/EtOAc 75:25 (1 L), 65:35 (1 L), 60:40 (2 L), 55:45 (2.5 L). At the end of 75:25 gradient, a yellow fraction elutes, and during the 60:40 elution a dark blue fraction emerges (NOTE: Neem powder from other brands sometimes does not contain this blue pigment). Nimbolide elutes at the tail end of the blue fraction. After removal of the solvent *in vacuo*, crude nimbolide is obtained as a dark green solid or thick oil, the TLC analysis of which is shown in SI-Figure 2D.

**A** **B** **C** **D**

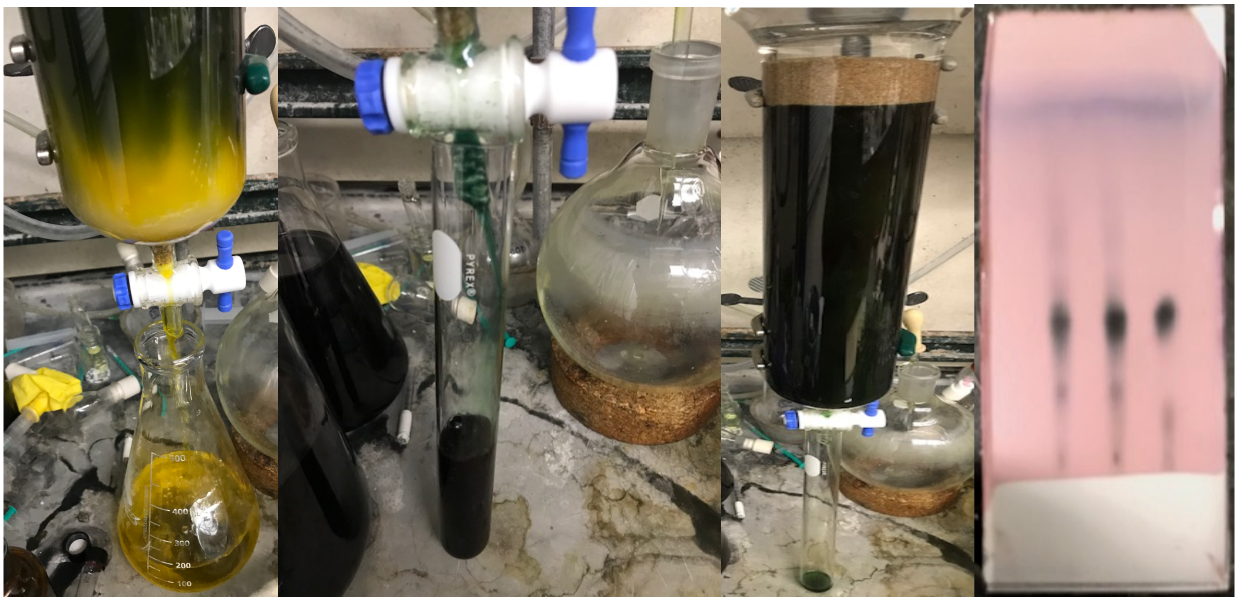

**SI-Figure 2.** Initial chromatographic event. **A**) Elution of the yellow fraction. **B**) elution of the blue fraction **C**) final column color (dark green). **D**) Thin layer chromatographic analysis (50:50 EtOAc/Hexane, *p*-anisaldehyde stain) of the combined nimbolide-containing fractions. Pure nimbolide is shown in the far-right lane, the column purified material in the left lane, and a co-spot in the center lane.

**Second Column Chromatography**: A column of silica gel (7 cm of diameter X 20 cm height) was packed and flushed with $CH_2Cl_2/Et_3N$ (100:1 v/v). Next, a solution of the green residue obtained after the 1st column in 25 mL of $CH_2Cl_2$ was loaded on the top. The column was eluted with the following gradient: $CH_2Cl_2$ (800 mL) $CH_2Cl_2$/acetone 60:1 (600 mL), 50:1 (1 L), 45:1 (2 L), 40:1 (3 L). Note that the green pigment should not move during this chromatographic event. After removal of solvent *in vacuo*, nimbolide is obtained as a dark brown solid, the TLC analysis of which is shown in SI-Figure 3C.

**A** **B** **C**

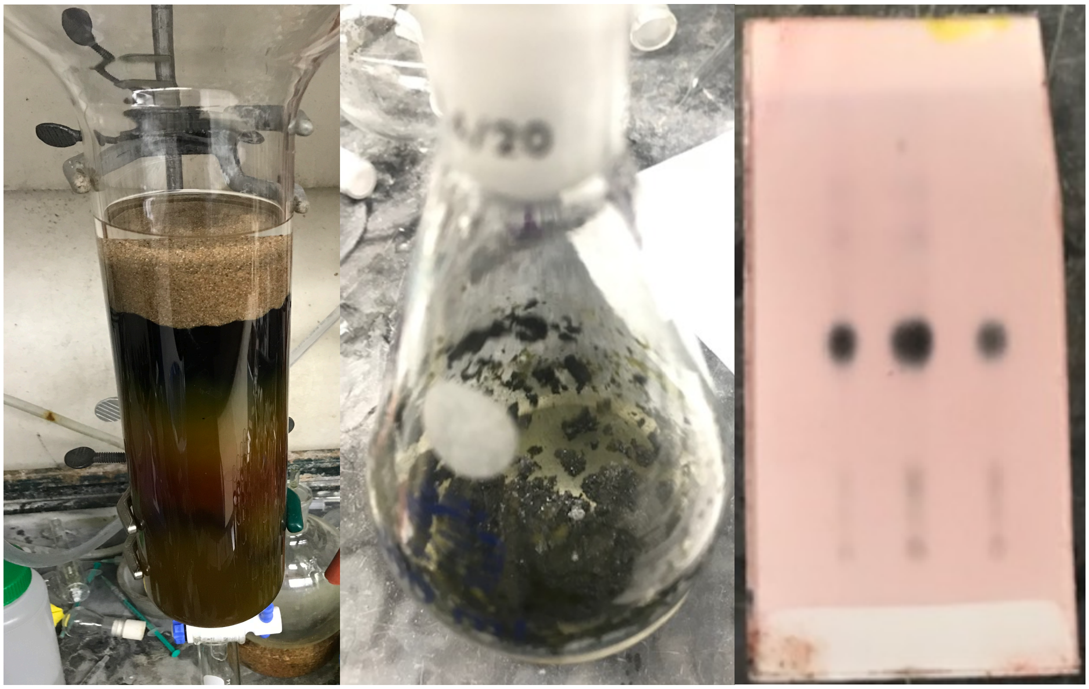

**SI-Figure 3.** Second chromatographic event. **A**) Column at the end of the elution. **B**) nimbolide obtained. **C**) Thin layer chromatographic analysis (50:50 EtOAc/Hexane, *p*-anisaldehyde stain) of the crude residue. Pure nimbolide is shown in the far-right lane, the column purified material in the left lane, and a co-spot in the center lane.

**Trituration to obtain analytically pure nimbolide.** The dark brown solid obtained after the 2nd column was diluted with 3 mL of $CH_2Cl_2$, followed by the addition of 20 mL of pentane. A brown solid precipitated out of solution and the liquid phase was removed by pipette. The solid was triturated 2 additional times in the same way (3 mL of $CH_2Cl_2$ then 20 mL of pentane) to furnish nimbolide as a white solid. A range of 1.1 g – 1.3 g of pure nimbolide was isolated using this procedure. No attempts were made to further purify the mother liquor which still contains significant quantities of nimbolide.

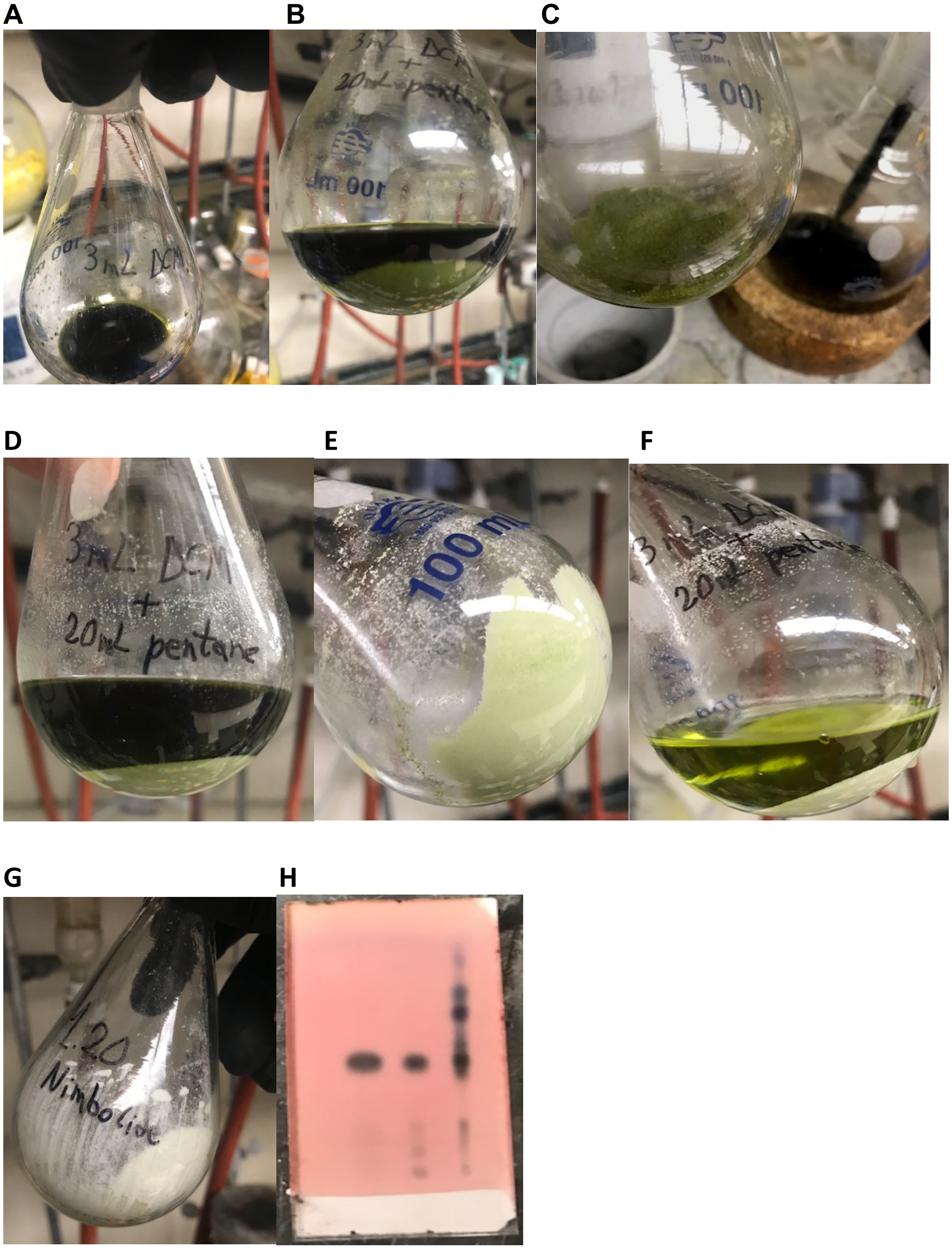


**SI-Figure 4.** Final trituration process. **A**) nimbolide in 3 ml of DCM. **B**) solid formed after addition of 20 ml of pentane. **C**) Removal of solvent from first trituration. **D-F**) final two triturations. **G**) Final pure nimbolide obtained. **H**) Thin layer chromatographic analysis (50:50 EtOAc/Hexane, *p*-anisaldehyde stain) of the trituration process.

The final, pure nimbolide obtained is shown in the far-left lane, an authentic standard in the middle lane, and the mother liquor is shown in the far-right lane.

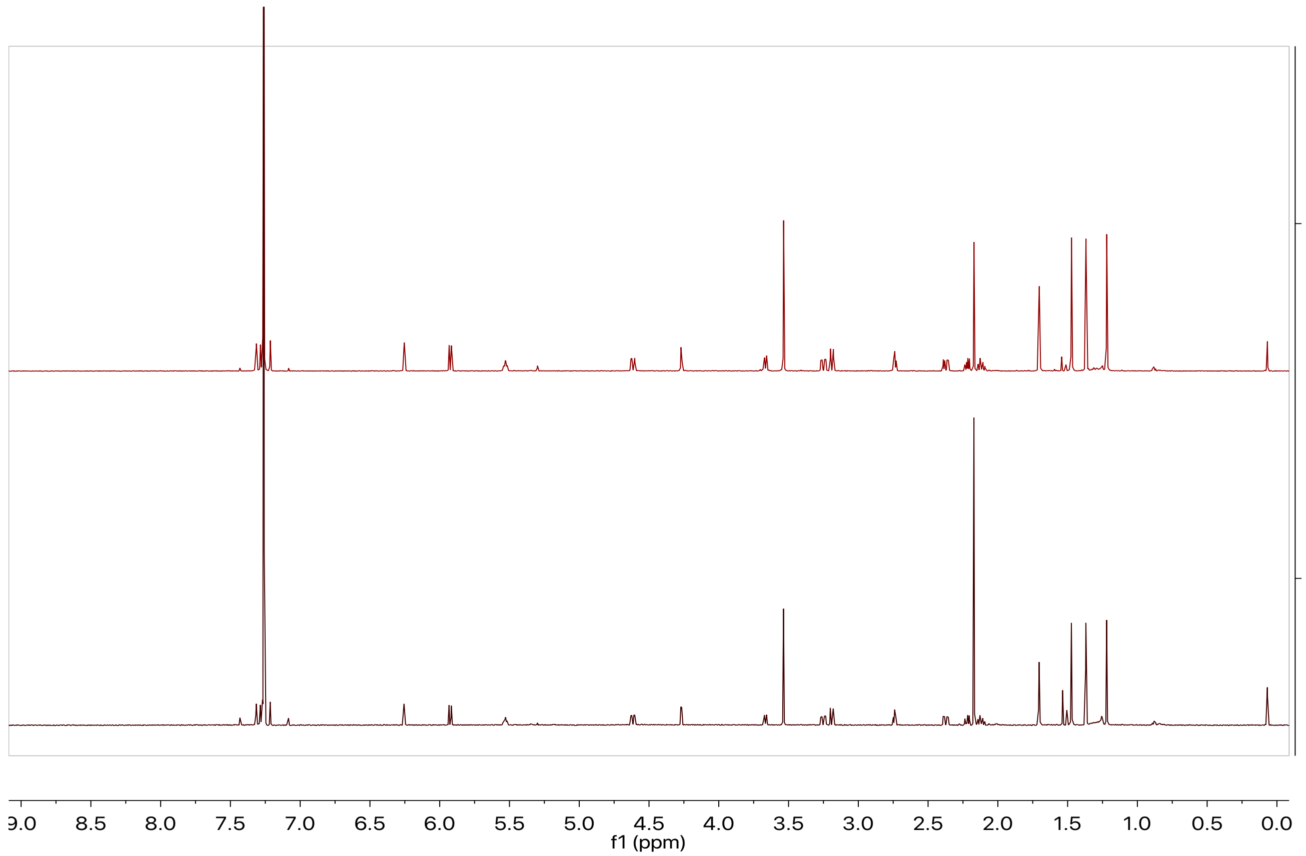


**SI-Figure 5.** 600 MHz $^1$H NMR Comparison of nimbolide in $CDCl_3$. (**Top**) Material obtained using the extraction protocol reported herein. (**Bottom**) Commercial nimbolide obtained from Sigma. (NOTE: nimbolide is quite acid sensitive and the $CDCl_3$ used was stored over $K_2CO_3$ prior to use).

**Synthesis and Characterization of Nimbolide-Dasatinib degraders BT1 and BT2**

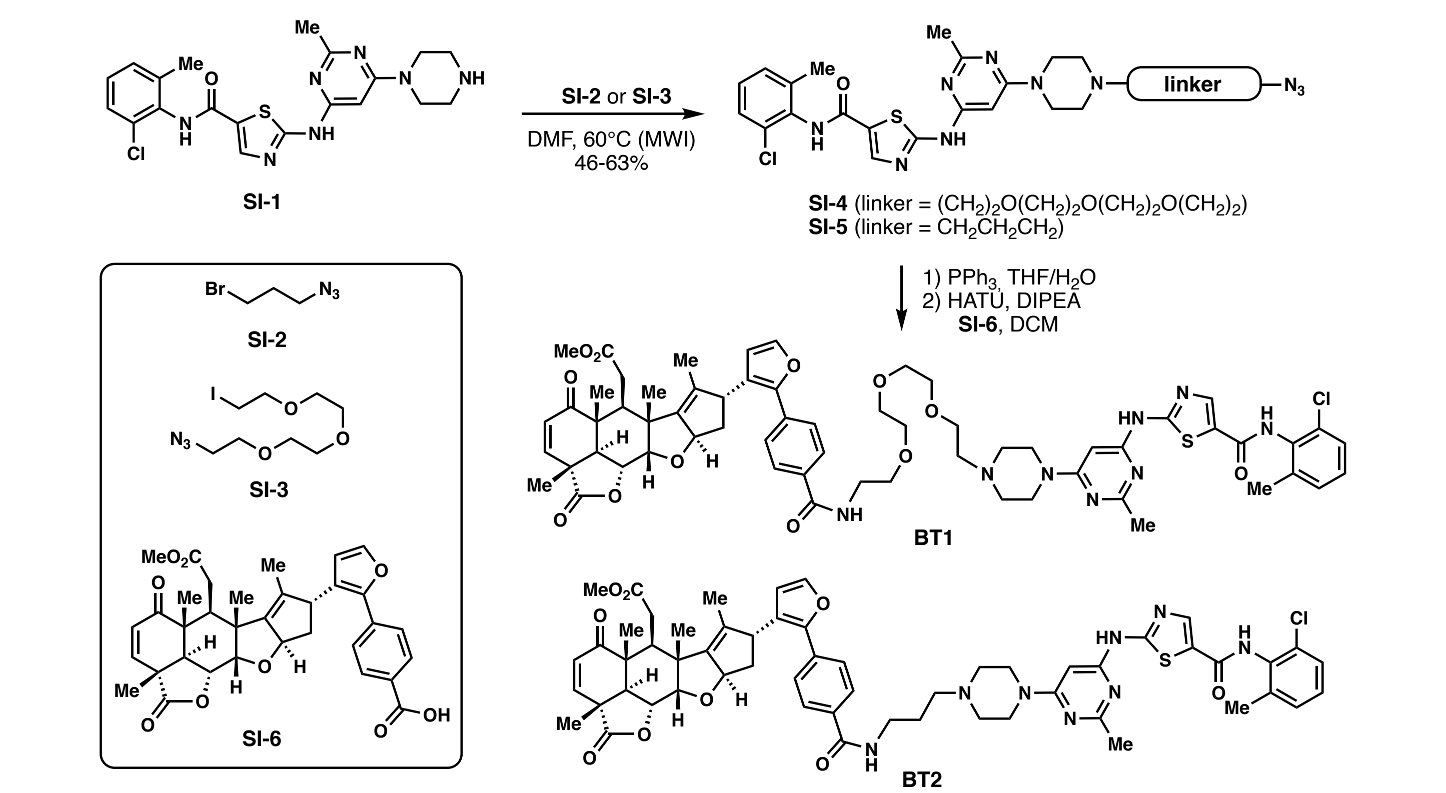


**Scheme SI-1.** Synthesis of nimbolide-dasatinib based degraders BT1 and BT2.

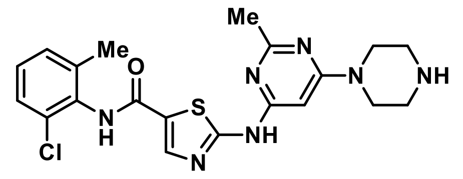

**Compound SI-1**: This compound was synthesized according to the procedure reported by Veach and coworkers.[1] (*J. Med. Chem.* **2007**, *50*, 5853-5857)

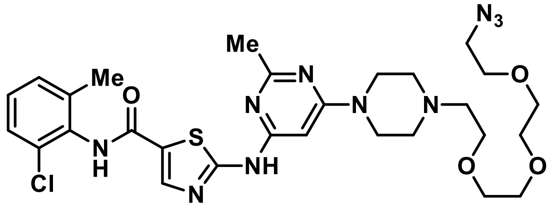

**Compound SI-4**: A 5 mL microwave tube equipped with a magnetic stir bar was charged with compound **SI-1** (111 mg, 0.25 mmol, 1 equiv), DMF (1.5 mL), DIPEA (65.0 µL, 0.375 mmol, 1.5 equiv) and iodide **SI-3** (51 µL, 0.25 mmol, 1.0 equiv). The tube was sealed and placed in a microwave reactor. The microwave reaction was performed at 60 °C for 1 h. After cooling to room temperature, the reaction mixture was transferred to a 50 mL round-bottom flask and the volatiles were remove under high vacuum. The remaining white solid was purified by column chromatography ($CH_2Cl_2$:MeOH = 95:5 to 93:7) to afford **SI-4** (101 mg, 63% yield) as a white solid: $^1H$ NMR (600 MHz, $CD_3OD$) δ 8.15 (s, 1H), 7.35 (d, *J* = 7.2 Hz, 1H), 7.30 – 7.18 (m, 2H), 6.02 (s, 1H), 3.72 – 3.62 (m, 16H), 3.37 (t, *J* = 4.8 Hz, 2H), 2.70 (t, *J* = 5.4 Hz, 2H), 2.68 (br, 4H), 2.48 (s, 3H), 2.32 (s, 3H); $^{13}C$ NMR (151 MHz, $CD_3OD$) δ 167.5, 165.3, 164.5, 163.3, 158.6, 142.2, 140.4, 134.4, 134.3, 130.1, 129.5, 128.3, 126.8, 83.9, 71.7, 71.7, 71.6, 71.4, 71.2, 69.4, 58.7, 54.1, 51.8, 44.7, 25.6, 18.7; IR (thin film) vmax 3182, 3061, 2925, 2870, 2094, 1718, 1616, 1536, 1189, 1144, 1010, 952, 781 $cm^{-1}$; HRMS (ESI) *calcd.* for $[C_{28}H_{38}O_4N_{10}ClS]^+$ ($[M+H]^+$) *m/z* 645.2481, found: 645.2476.

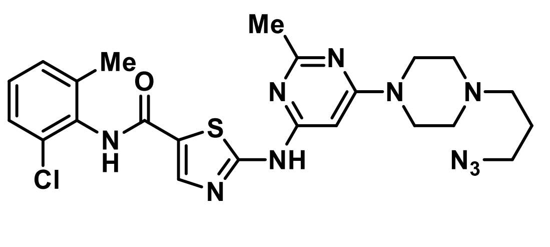

**Compound SI-5**: A 30 mL microwave tube equipped with a magnetic stir bar was charged with compound **SI-1** (533 mg, 1.20 mmol, 1 equiv), DMF (12 mL), DIPEA (0.418 mL, 2.4 mmol, 2 equiv) and bromide **SI-2** (0.178 mL, 1.44 mmol, 1.2 equiv). The tube was sealed and placed in the microwave reactor. The microwave reaction was performed at 60 °C for 1.5 h. After cooling to room temperature, the reaction mixture was transferred into a 100 mL round-bottom flask and the DMF and other volatiles were removed *in vacuo*. The resulting white solid was then suspended in $CH_2Cl_2$/MeOH (90:10 v/v), passed through a short column of silica gel and eluted with $CH_2Cl_2$/MeOH (90:10). The resulting solution was concentrated *in vacuo* to afford the crude product, which was further purified by column chromatography ($CH_2Cl_2$:MeOH = 95:5) to yield **SI-5** (290 mg, 46% yield) as a white solid: $^1$H NMR (600 MHz, $(CD_3)_2SO$) δ 11.47 (s, 1H), 9.88 (s, 1H), 8.22 (s, 1H), 7.40 (d, *J* = 7.7 Hz, 1H), 7.33 – 7.22 (m, 2H), 6.05 (s, 1H), 3.51 (br, 4H), 3.39 (t, *J* = 6.7 Hz, 2H), 2.43 (br, 4H), 2.40 (s, 3H), 2.38 (t, *J* = 6.9 Hz, 2H), 2.24 (s, 3H), 1.73 (p, *J* = 6.8 Hz, 2H); $^{13}$C NMR (151 MHz, $(CD_3)_2SO$) δ 165.2, 162.6, 162.4, 159.9, 156.9, 140.8, 138.8, 133.5, 132.4, 129.0, 128.1, 127.0, 125.7, 82.6, 54.6, 52.2, 48.9, 43.6, 25.61, 25.58, 18.3; IR (thin film) vmax 3205, 3060, 2938, 2847, 2093, 1715, 1621, 1536, 1182, 1142, 998, 862 $cm^{-1}$; HRMS (ESI) *calcd* for $[C_{23}H_{28}ON_{10}ClS]^+$ ($[M+H]^+$): *m/z* 527.1851, found: 527.1850.

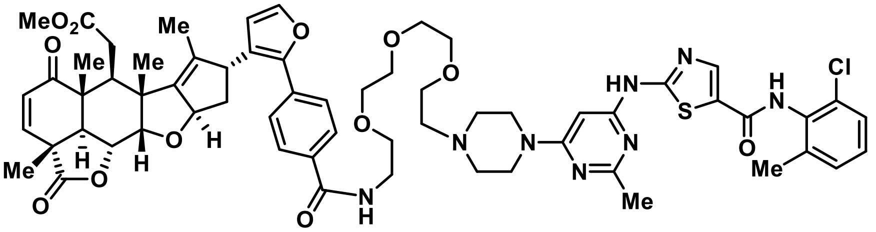

**Degrader BT1**: *i.* To a 10 mL reaction tube was added **SI-4** (10.6 mg, 0.0164 mmol, 1.0 equiv), triphenylphosphine (8.6 mg, 0.033 mmol, 2.0 equiv), THF (0.6 mL) and $H_2O$ (0.06 mL). The resulting solution was heated in a 50 °C oil bath for 12 hours. After full consumption of the starting material was indicated by TLC, all the volatiles in the reaction mixture was removed by rotary evaporation at 60 °C. The resulting crude pale-yellow solid containing the primary amine was further dried under high vacuum for 12 hours before being subjected to the next step.

*ii*. To a separate 10 mL reaction tube was added acid **SI-6** (0.016 mmol, 1.0 equiv),[2] HATU (6.1 mg, 0.016 mmol, 1.0 equiv), DIPEA (5.6 µL, 0.032 mmol, 2.0 equiv) and $CH_2Cl_2$ (0.4 mL). After stirring at room temperature for 10 hours, the reaction was quenched with saturated *aq*. $NH_4Cl$ and extracted with $CH_2Cl_2$ (3 × 1 mL). The combined organic layer was dried over $Na_2SO_4$ and concentrated *in vacuo*. The resulting activated ester was then transferred into the reaction tube containing the crude primary amine, DIPEA (5.6 µL, 0.032 mmol, 2.0 equiv) and $CH_2Cl_2$ (0.4 mL). The reaction mixture was allowed to stir at room temperature for 12 hours before quenching with saturated aq. $NH_4Cl$. The layers were separated, and the aqueous phase was extracted with $CH_2Cl_2$ (3×1 mL). The combined organic layer was dried over $Na_2SO_4$ and concentrated *in vacuo*. The residue was purified by column chromatography (7% MeOH/$CH_2Cl_2$ with 0.2% $Et_3N$) to afford degrader **BT1** (10.5 mg, 54% yield) as a colorless oil which slowly solidifies. $^1$H NMR (600 MHz, $(CD_3)_2CO$) δ 10.38 (br, 1H), 9.03 (br, 1H), 8.20 (s, 1H), 8.03 (d, *J* = 8.4 Hz, 2H), 7.83 (br, 1H), 7.67 (d, *J* = 8.4 Hz, 2H), 7.56 (d, *J* = 1.8 Hz, 1H), 7.34 (d, *J* = 7.7 Hz, 1H), 7.30 (d, *J* = 9.7 Hz, 1H), 7.28 – 7.20 (m, 2H), 6.45 (d, *J* = 1.8 Hz, 1H), 6.17 (s, 1H), 5.88 (d, *J* = 9.7 Hz, 1H), 5.51 (ddd, *J* = 8.6, 3.6, 1.8 Hz, 1H), 5.00 (dd, *J* = 12.4, 3.7 Hz, 1H), 4.28 (d, *J* = 3.6 Hz, 1H), 4.20 (d, *J* = 9.0 Hz, 1H), 3.68 (t, *J* = 5.7 Hz, 2H), 3.64 – 3.55 (m, 16H), 3.62 (s, 3H), 3.26 (dd, *J* = 16.7, 5.4 Hz, 1H), 3.12 (d, *J* = 12.5 Hz, 1H), 2.78 (t, *J* = 5.4 Hz, 1H), 2.55 (br, 6H), 2.48 (dd, *J* = 16.7, 5.4 Hz, 1H), 2.44 (s, 3H), 2.33 (s, 3H), 2.24 (ddd, *J* = 12.0, 8.6, 8.4 Hz, 1H), 2.18 (dd, *J* = 12.0, 6.9 Hz, 1H), 1.68 (d, *J* = 1.5 Hz, 3H), 1.50 (s, 3H), 1.44 (s, 3H), 1.30 (s, 3H); $^{13}$C NMR (151 MHz, $(CD_3)_2CO$) δ 201.6, 176.1, 173.9, 167.1, 166.6, 163.9, 160.9, 158.2, 150.7, 149.0, 147.4, 143.3, 141.5, 140.1, 136.4, 134.7, 134.6, 134.2, 133.5, 131.5, 129.8, 128.9, 128.7, 128.5, 127.8, 127.4, 126.4, 125.6, 113.23, 88.7, 84.0, 83.4, 74.1, 71.3, 71.2, 71.08, 71.06, 70.3, 70.1, 58.5, 54.0, 51.9, 51.50, 50.5, 49.1, 46.3, 44.8, 44.7, 41.92, 41.90, 40.6, 32.8, 25.9, 18.9, 18.7, 17.20, 15.2, 13.4; IR (thin film) vmax 3385, 2923. 2852, 1733, 1611,1563, 1449, 1417, 1290, 1197, 1117, 824, 714 $cm^{-1}$; HRMS (ESI) *calcd* for $[C_{62}H_{72}O_{12}N_8ClS]^+$ ($[M+H]^+$): *m/z* 1187.4673, found: 1187.4663.

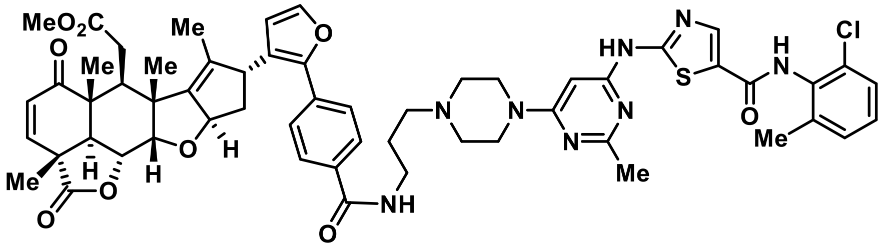

**Degrader BT2**: *i.* To a 10 mL reaction tube was added **SI-5** (14.8 mg, 0.0281 mmol, 1.0 equiv), triphenylphosphine (14.7 mg, 0.056 mmol, 2.0 equiv), THF (1.0 mL) and $H_2O$ (0.1 mL). The resulting solution was heated in a 50 °C oil bath for 12 hours. After fully consumption of the starting material was indicated by TLC, all the volatiles in the reaction mixture was removed by rotary evaporation at 60 °C. The resulting crude pale-yellow solid containing the primary amine was further dried under high vacuum overnight before being subjected to the next step.

*ii.* To another 10 mL reaction tube was added acid **SI-6** (0.0261 mmol, 0.93 equiv), HATU (9.6 mg, 0.025 mmol, 0.9 equiv), DIPEA (9.8 μL) and $CH_2Cl_2$ (0.6 mL). After stirring at room temperature for 1 hour, this reaction mixture was transferred via syringe into the reaction tube containing the crude primary amine and $CH_2Cl_2$ (0.2 mL). The resulting mixture was stirred for 12 hours before quenching with saturated aq. $NH_4Cl$. The layers were separated, and the aqueous phase was extracted with $CH_2Cl_2$ (3×1 mL). The combined organic layer was dried over $Na_2SO_4$ and concentrated *in vacuo*. The residue was purified by column chromatography (5% MeOH/$CH_2Cl_2$ with 0.2% $Et_3N$), followed by HPLC to afford degrader **BT2** (7.8 mg, 26% yield) as a colorless oil which slowly solidifies. $^1$H NMR (600 MHz, $(CD_3)_2CO$) δ 9.06 (br, 1H), 8.25 (br, 1H), 8.20 (br, 1H), 7.97 (d, *J* = 8.4 Hz, 2H), 7.66 (d, *J* = 8.4 Hz, 2H), 7.55 (d, *J* = 1.7 Hz, 1H), 7.34 (d, *J* = 7.6 Hz, 1H), 7.30 (d, *J* = 9.7 Hz, 1H), 7.28 – 7.20 (m, 2H), 6.43 (d, *J* = 1.8 Hz, 1H), 6.18 (s, 1H), 5.87 (d, *J* = 9.7 Hz, 1H), 5.49 (t, *J* = 7.2 Hz, 1H), 4.98 (dd, *J* = 12.5, 3.6 Hz, 1H), 4.26 (d, *J* = 3.6 Hz, 1H), 4.14 (t, *J* = 4.8 Hz, 1H), 3.67 (br, 4H), 3.61 (s, 3H), 3.58 – 3.47 (m, 2H), 3.24 (dd, *J* = 16.5, 5.1 Hz, 1H), 3.10 (d, *J* = 12.4 Hz, 1H), 2.76 (t, *J* = 5.4 Hz, 1H), 2.63 (br, 6H), 2.46 (s, 3H), 2.45 (dd, *J* = 16.5, 5.5 Hz, 1H), 2.33 (s, 3H), 2.17 (dd, *J* = 7.4, 5.1 Hz, 2H), 1.88 (quint, *J* = 6.0 Hz, 2H), 1.63 (d, *J* = 1.3 Hz, 3H), 1.50 (s, 3H), 1.39 (s, 3H), 1.28 (s, 3H); $^{13}$C NMR (151 MHz, $(CD_3)_2CO$) δ 201.6, 176.1, 173.9, 166.8, 166.7, 164.0, 160.9, 158.2, 150.8, 149.0, 147.3, 143.3, 141.4, 140.1, 136.4, 134.62, 134.59, 134.52, 133.5, 131.5, 129.8, 128.9, 128.5, 128.5, 127.8, 127.5, 126.4, 125.6, 113.2, 88.7, 83.9, 83.5, 74.1, 57.6, 53.6, 51.9, 51.5, 50.5, 49.1, 46.2, 44.7, 44.6, 41.91, 41.88, 39.8, 32.8, 26.5, 25.9, 18.9, 18.7, 17.2, 15.3, 13.3; IR (thin film) $\nu_{max}$ 3252, 2923, 2853, 1781, 1735, 1611, 1577, 1508, 1463, 1416, 1290, 1198, 851, 750 cm$^{-1}$; HRMS (ESI) *calcd.* for $[C_{57}H_{62}O_9N_8ClS]^+$ ($[M+H]^+$): *m/z* 1069.4044, found: 1069.4047.

### References


1. Veach, D. R.; Namavari, M.; Pillarsetty, N.; Santos, E. B.; Tatiana Beresten-Kochetkov, T.; Lambek, C.; Punzalan, B. J.; Antczak, C.; Smith-Jones, P. M.; Djaballah, H.; Clarkson, B.; Larson, S. M. *J. Med. Chem.* **2007**, *50*, 5853-5857
2. Spradlin, J. N.; Hu, X.; Ward, C. C.; Brittain, S. M.; Jones, M. D.; Ou, L.; To, M.; Proudfoot, A.; Ornelas, E.; Woldegiorgis, M.; Olzmann, J. A. Bussiere, D. E.; Thomas, J. R.; Tallarico, J. A.; McKenna, J. M.; Schirle, M.; Maimone, T. J.; Nomura, D. K. *Nat. Chem. Biol.* **2019**, *15*, 747-755

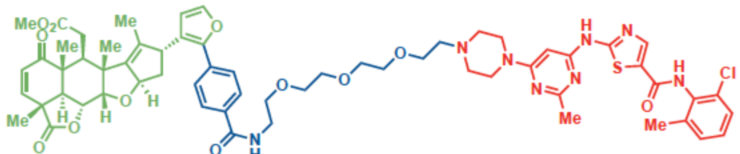

BT1

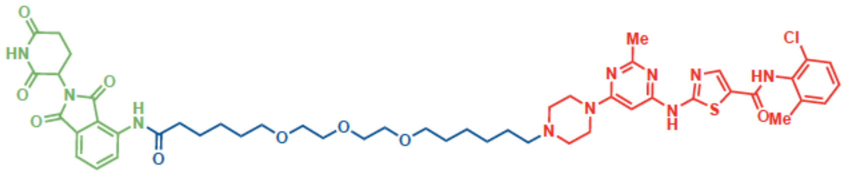

CRBN-DAS

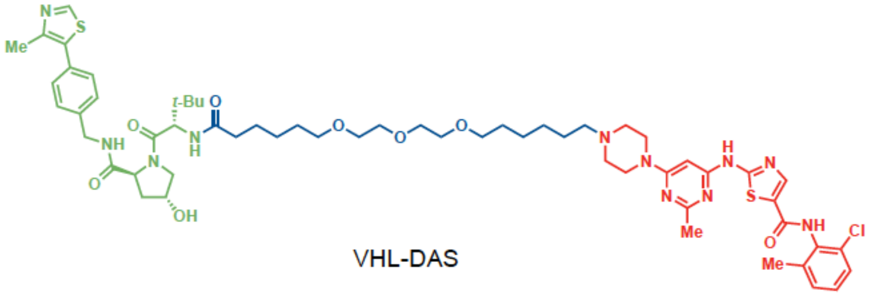

VHL-DAS

**Figure S1.** Structures of BT1, CRBN-dasatinib, and VHL-dasatinib. This figure is related to **Figure 3.**
